# Ovariectomy-Induced Arterial Stiffening Differs from Vascular Aging and is Reversed by GPER Activation

**DOI:** 10.1101/2023.08.10.552881

**Authors:** Isabella M. Kilanowski-Doroh, Alexandra B. McNally, Tristen Wong, Bruna Visniauskas, Sophia A. Blessinger, Ariane Imulinde Sugi, Chase Richard, Zaidmara Diaz, Alec Horton, Christopher A. Natale, Benard O. Ogola, Sarah H. Lindsey

## Abstract

Arterial stiffness is a cardiovascular risk factor and dramatically increases as women transition through menopause. The current study assessed whether a mouse model of menopause increases arterial stiffness in a similar manner to aging, and whether activation of the G protein-coupled estrogen receptor (GPER) could reverse stiffness. Female C57Bl/6J mice were ovariectomized (OVX) at 10 weeks of age or aged to 52 weeks, and some mice were treated with GPER agonists. OVX and aging increased pulse wave velocity to a similar extent independent of changes in blood pressure. Aging increased carotid wall thickness, while OVX increased material stiffness without altering vascular geometry. RNA-Seq analysis revealed that OVX downregulated smooth muscle contractile genes. The enantiomerically pure GPER agonist, LNS8801, reversed stiffness in OVX mice to a greater degree than the racemic agonist G-1. In summary, OVX and aging induced arterial stiffening via potentially different mechanisms. Aging was associated with inward remodeling while OVX induced material stiffness independent of geometry and a loss of the contractile phenotype. This study helps to further our understanding of the impact of menopause on vascular health and identifies LNS8801 as a potential therapy to counteract this detrimental process in women.

## INTRODUCTION

Pulse wave velocity (PWV) is a well-documented measurement of arterial stiffness, and elevated PWV is an independent determinant of cardiovascular outcomes (1, 2). While aging-induced arterial stiffness is expected due to regeneration and remodeling across the lifespan, acute injury and chronic inflammatory conditions such as hypertension also contribute (3, 4). Arterial stiffening is often associated with extracellular matrix remodeling such as elastin fragmentation and increased collagen deposition (5), but these mechanisms do not fully explain the process behind vascular stiffening (6, 7). The contribution of vascular smooth muscle cells and cell-matrix interactions are less studied but play a significant role in models of aging and hypertension (8–10).

Since aging is a major contributor, the prevalence of arterial stiffening will increase as life expectancy continues to extend (11). Despite this increased risk, arterial stiffness is not routinely measured or treated clinically (12). Initial work in this field suggested that hypertension preceded arterial stiffness, but recent studies show that the relationship is bidirectional since older individuals with optimal blood pressure and no comorbidities exhibit high PWV (13, 14). Several mechanisms contribute to this phenomenon of “vascular aging”, including adiposity, inflammation, and endothelial dysfunction (15, 16). Because arterial compliance is similar across sexes at birth but diverges after puberty (17), sex and sex hormones also play a role in arterial stiffening. Recent data from the Study of Women Across the Nation (SWAN) shows arterial stiffness drastically increases in women within one year of the final menstrual period without any alterations in blood pressure, strongly indicating an essential role for estrogen in vascular homeostasis and remodeling (18).

Our group previously showed that PWV is higher in adult male versus female mice, but this sex difference is lost with hypertension or aging (19, 20). Our previous studies also have explored the role of membrane-initiated estrogen signaling in female protection from cardiovascular disease (21–23). We find that pharmacological activation of the G protein-coupled estrogen receptor (GPER) induces vasodilation, attenuates hypertension and salt-induced vascular remodeling, and promotes diastolic function (24–30). In contrast, genetic deletion of this receptor increases arterial stiffness only in female mice (31–33). These studies implicate a role for GPER in arterial stiffness, but the impact of estrogen loss was not assessed. Moreover, while the commercially available agonist G-1 contains multiple enantiomers, the novel GPER agonist LNS8801 is enantiomerically pure and is currently being tested in clinical trials for cancer (34, 35). Therefore, the current study hypothesized that estrogen loss via ovariectomy would recapitulate the impact of chronological aging on arterial stiffening, while pharmacological targeting of GPER would reverse the impact of estrogen loss.

## RESULTS

Neither OVX nor aging significantly impacted body weight, heart weight, or kidney weight (Figure 1A-C). Uterine weight was significantly lower in ovariectomized mice but higher in middle-aged mice compared with controls (Figure 1D). Both OVX and aging increased intracarotid PWV (icPWV) and aortic PWV (aPWV) to a similar extent (Figure 2A-2B). Systolic blood pressure as well as pulse pressure was similar across groups (Figure 2C-2D).

**Figure 1.**
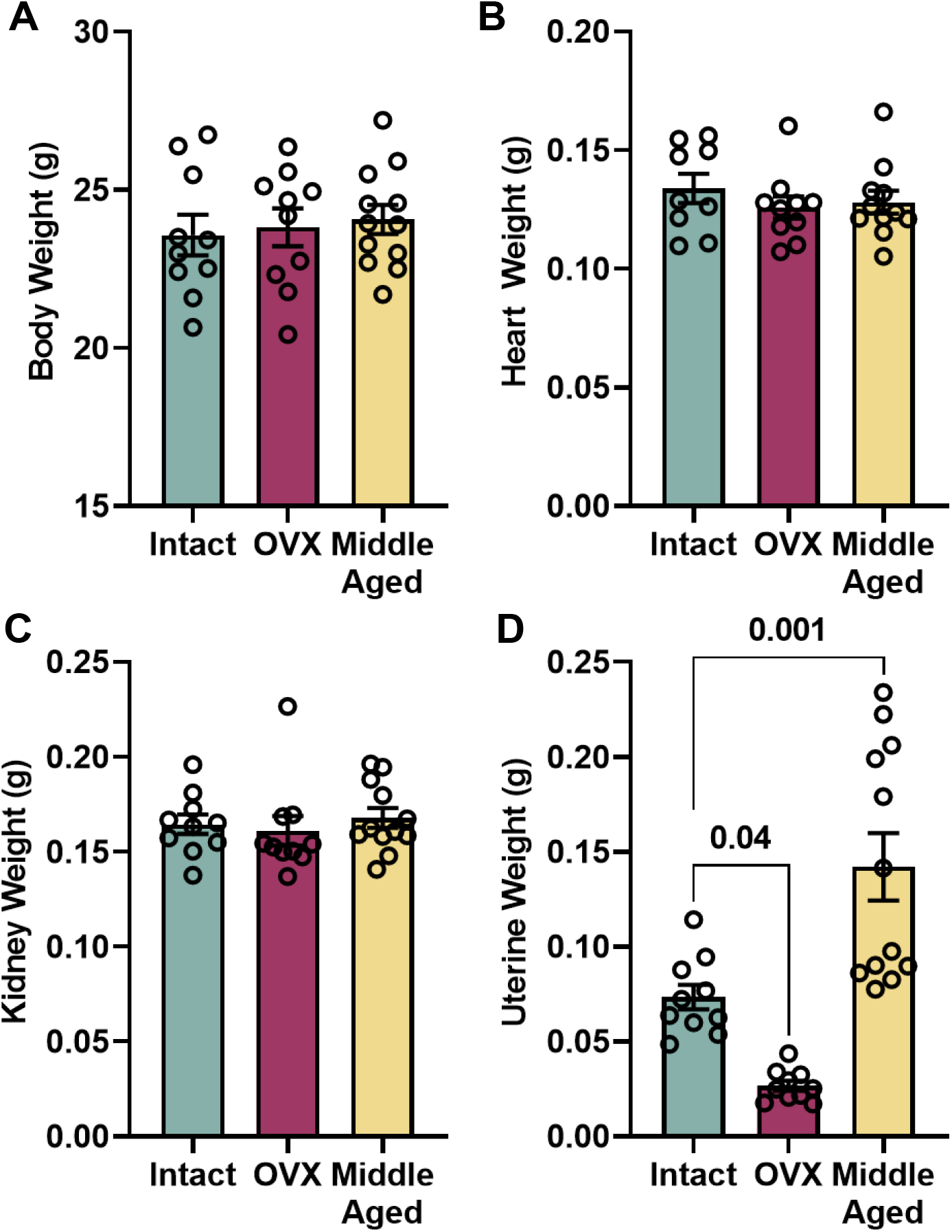
Impact of OVX and aging on body and tissue weights. Statistical analysis was performed using one-way ANOVA with Bonferroni’s multiple comparisons test shown on the graphs. Values are mean ± SEM, N=10-11 per group. No significant differences were seen in **(A)** body weight, P=0.82; **(B)** heart weight, P=0.55; and **(C)** kidney weight, P=0.73. **(D)** OVX significantly decreased while aging significantly increased uterine weight compared with intact controls, P<0.0001.

**Figure 2.**
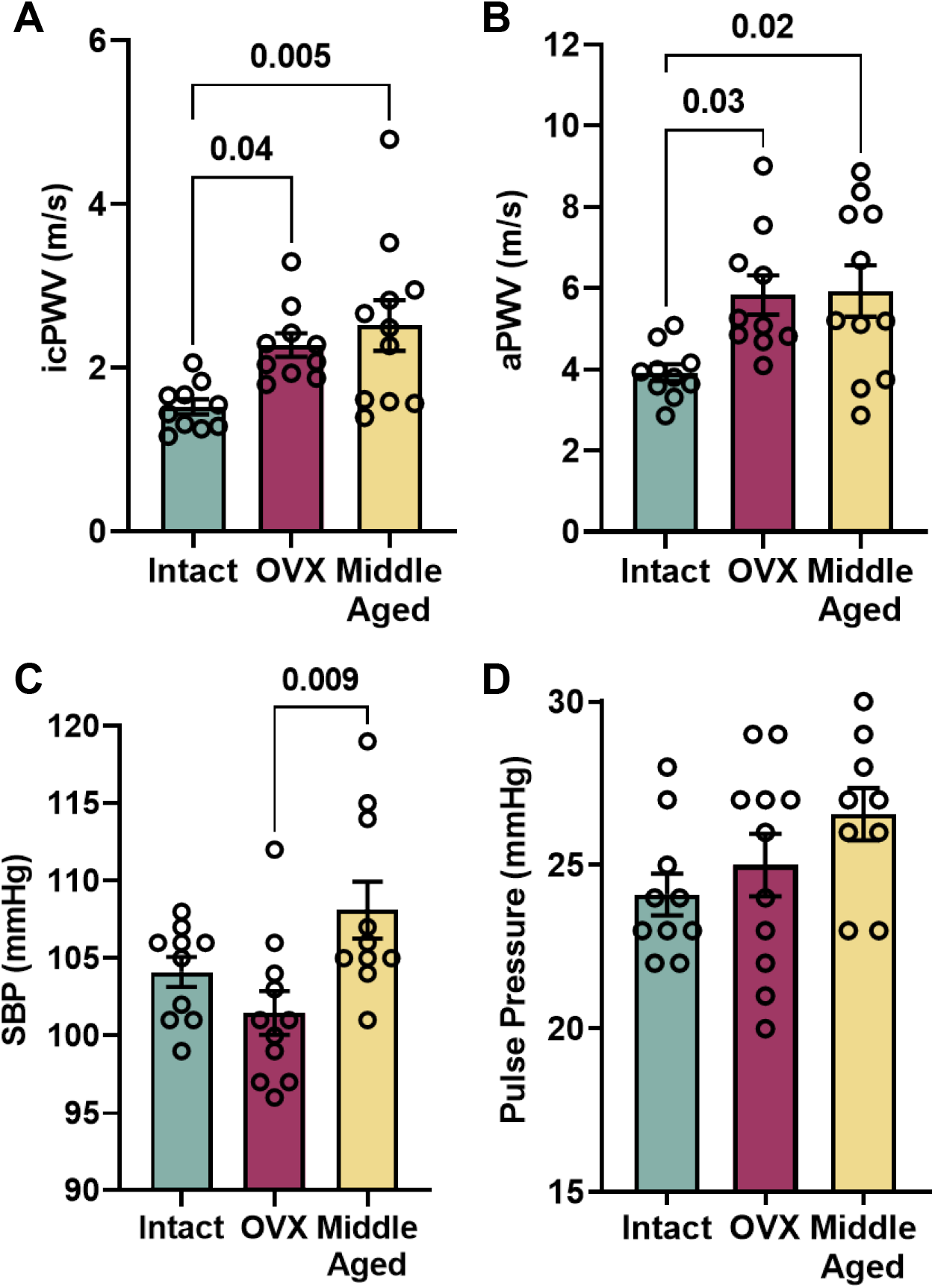
Impact of OVX and aging on blood pressure and PWV. Statistical analysis was performed using one-way ANOVA with Bonferroni’s multiple comparisons test shown on the graphs. Values are mean ± SEM, N=10-11 per group. **(A)** Intracarotid PWV (icPWV) and **(B)** Aortic PWV (aPWV) were significantly increased in both OVX and middle-aged groups compared with intact controls, P=0.007 and P=0.01. **(C)** Systolic blood pressure (SBP) was different by ANOVA, P=0.01, due to a difference between the OVX and middle-aged groups. **(D)** Pulse pressure was not different between groups, P=0.14.

Wall thickness and wall/lumen ratio were significantly greater only in the middle-aged group (Figure 3A-B). This change in geometry was due to a decrease in luminal diameter, with no change in external diameter (Figure 3C-D). Pressure myography showed that OVX carotids had greater distension between 60-110 mmHg (Figure 4A). The stress-strain curve was shifted leftward in OVX, indicating an increase in material stiffness, but was similar in middle-aged versus young intact mice (Figure 4B). Beta stiffness index (Figure 4C) and incremental modulus (Figure 4D) were both calculated between the physiological range of 80-120 mmHg, and again show that OVX but not aging increased material stiffness.

**Figure 3.**
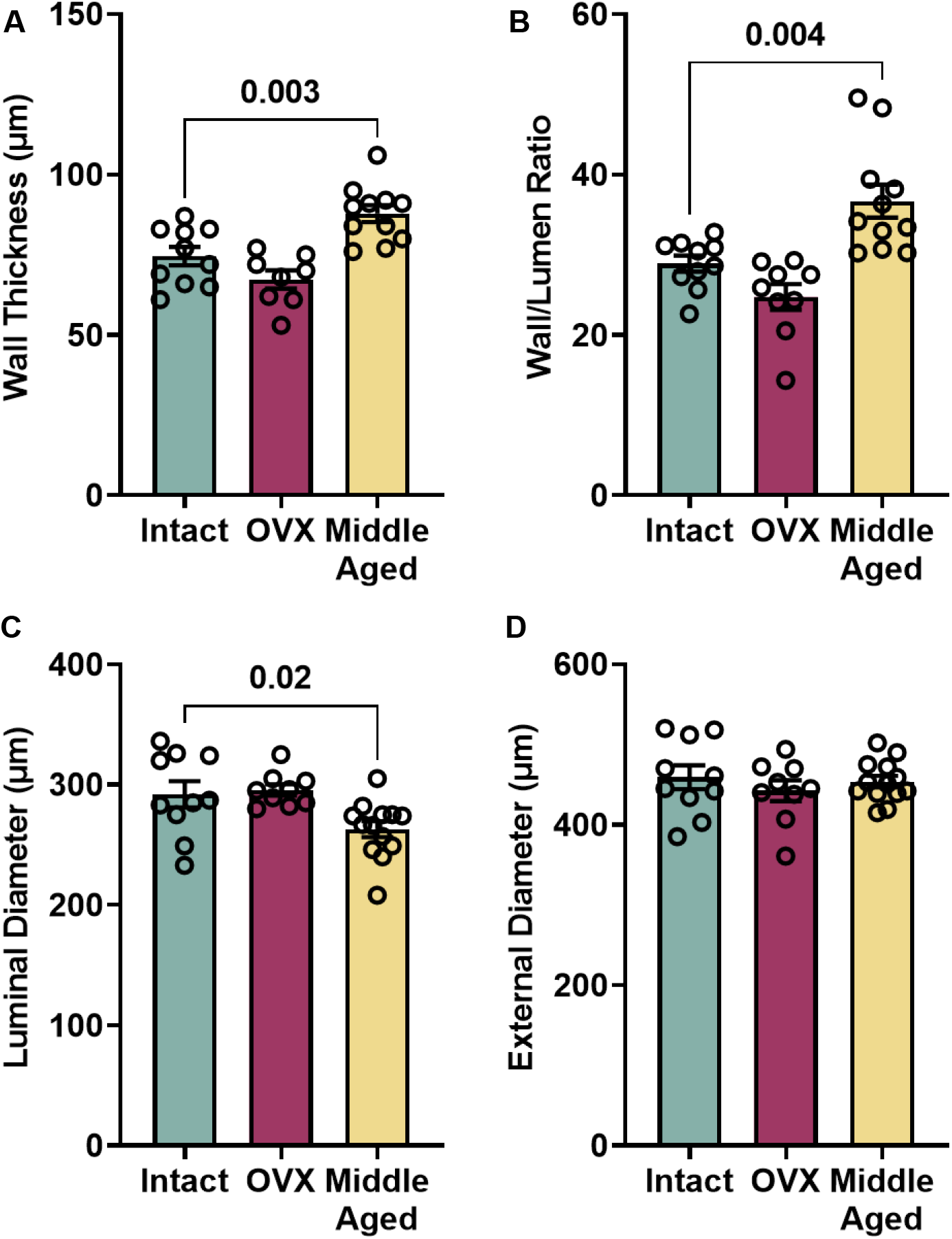
Impact of OVX and aging on vascular geometry. Statistical analysis was performed using one-way ANOVA with Bonferroni’s multiple comparisons test shown on the graphs. Values are mean ± SEM, N=10-11 per group. **(A)** Wall thickness and **(B)** wall-to-lumen ratio were significantly increased in middle-aged but not OVX mice, P<0.001. **(C)** Luminal diameter was significantly decreased in middle-aged mice, P=0.009. **(D)** External diameter was not different between groups, P=0.61.

**Figure 4.**
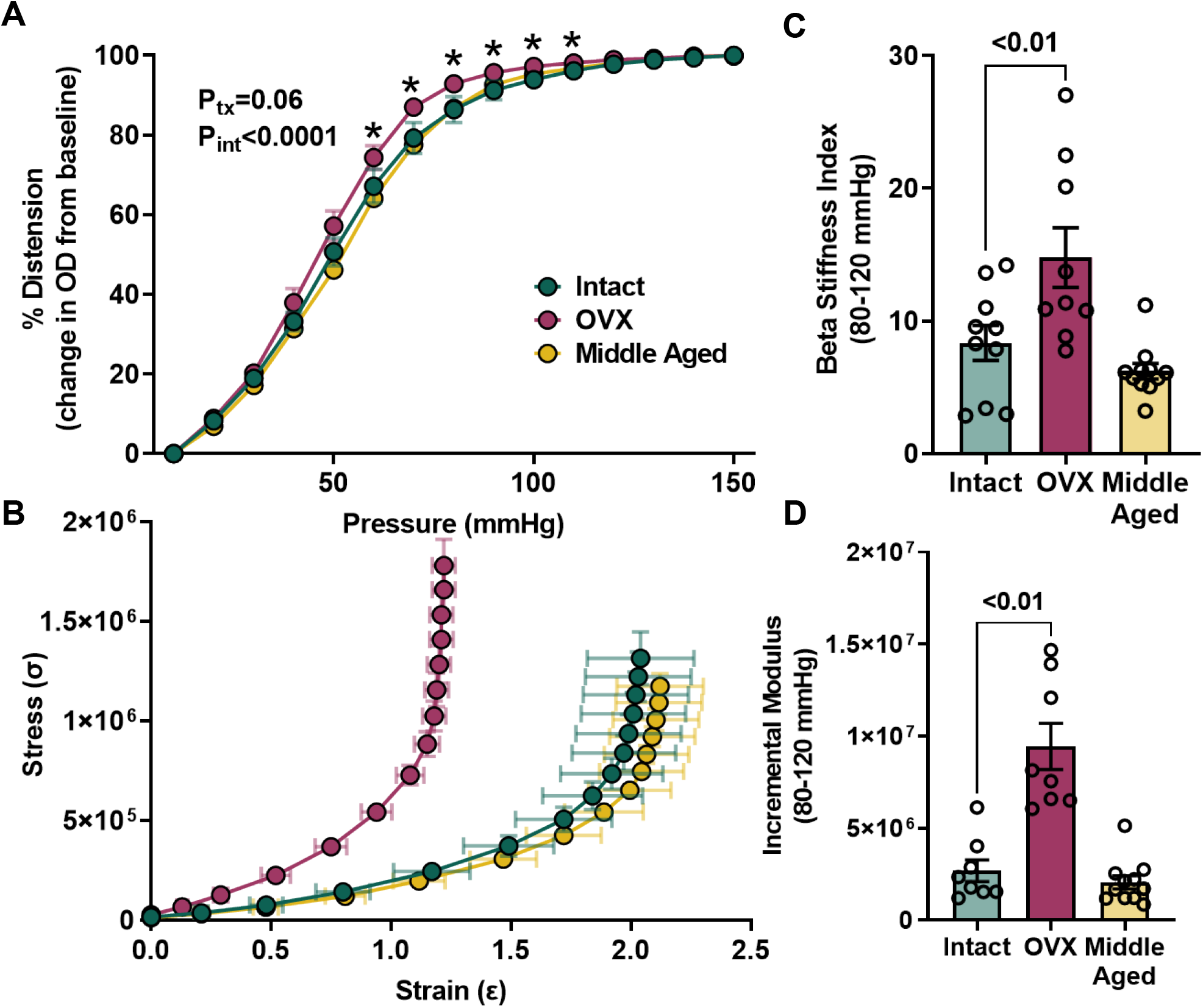
Impact of OVX and aging on material stiffness. Values are mean ± SEM, N=10-11 per group. **(A)** There was a significant interaction between pressure and treatment group (tx) due to greater carotid distensibility in OVX carotids over the range of physiological pressures. Two-way ANOVA and Bonferroni post-hoc results are shown on the graph. **(B)** A leftward shift of the OVX stress-strain curve indicates increased material stiffness. **(C)** Beta stiffness index and **(D)** incremental modulus calculated between the physiological range of 80-120 mmHg shows that OVX but not aging increases material stiffness. One-way ANOVA, P<0.01.

Analysis of aortic cross sections showed no significant differences in elastin content between groups (Figure 5A), while both smooth muscle and collagen content within the medial layer were increased only in the middle-aged group (Figure 5B-5C). Alcian blue staining for glycosaminoglycans (GAGs) showed a significant decrease in GAG content in OVX but not middle-aged aortas (Figure 5D). While histological analysis first focused on analysis of the medial layer, we noted changes in the size of the adventitial layer and thus quantified this in all groups using stained sections. In OVX aortas, the medial layer was significantly decreased while the adventitial layer was significantly increased (Figure 6A), resulting in no change in total area and confirming the similar external diameters obtained from pressure myography experiments (Figure 6B). In contrast, aging increased both the medial and adventitial layers to contribute to the overall increase in wall thickness. Despite the fact that OVX and aging had divergent effects on layer composition, both induced a strikingly similar decrease when expressed as the ratio of media to adventitia (Figure 6C).

**Figure 5.**
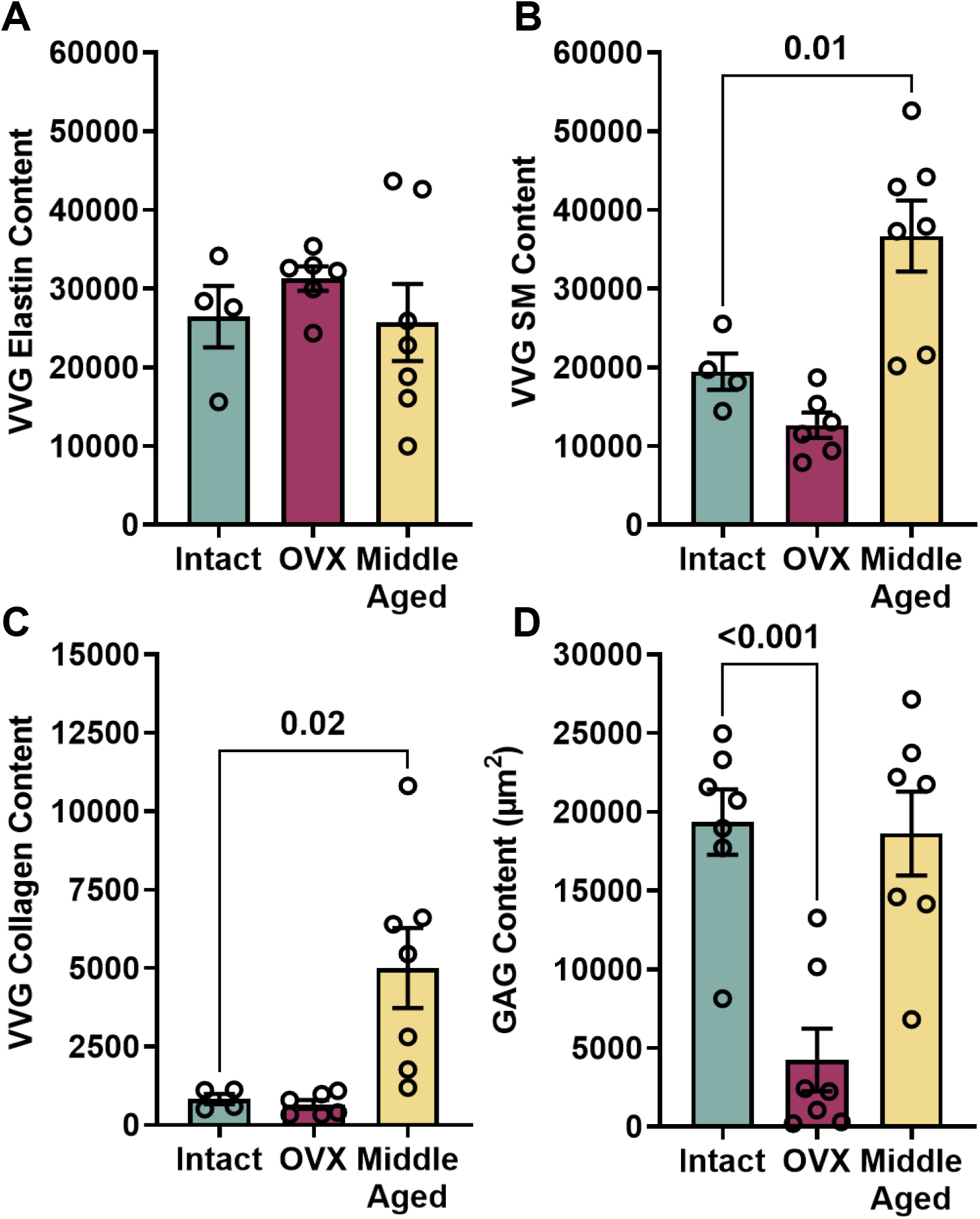
Aortic Histology. Statistical analysis was performed using one-way ANOVA with Bonferroni’s multiple comparisons test shown on the graphs. Values are mean ± SEM, N=10-11 per group. VVG stain showed no significant difference in **(A)** elastin, P=0.56, while **(B)** smooth muscle, P<0.001, and **(C)** collagen, P=0.006, were increased only in the middle-aged group. **(D)** OVX significantly decreased area fraction of glycosaminoglycans, P<0.001.

**Figure 6.**
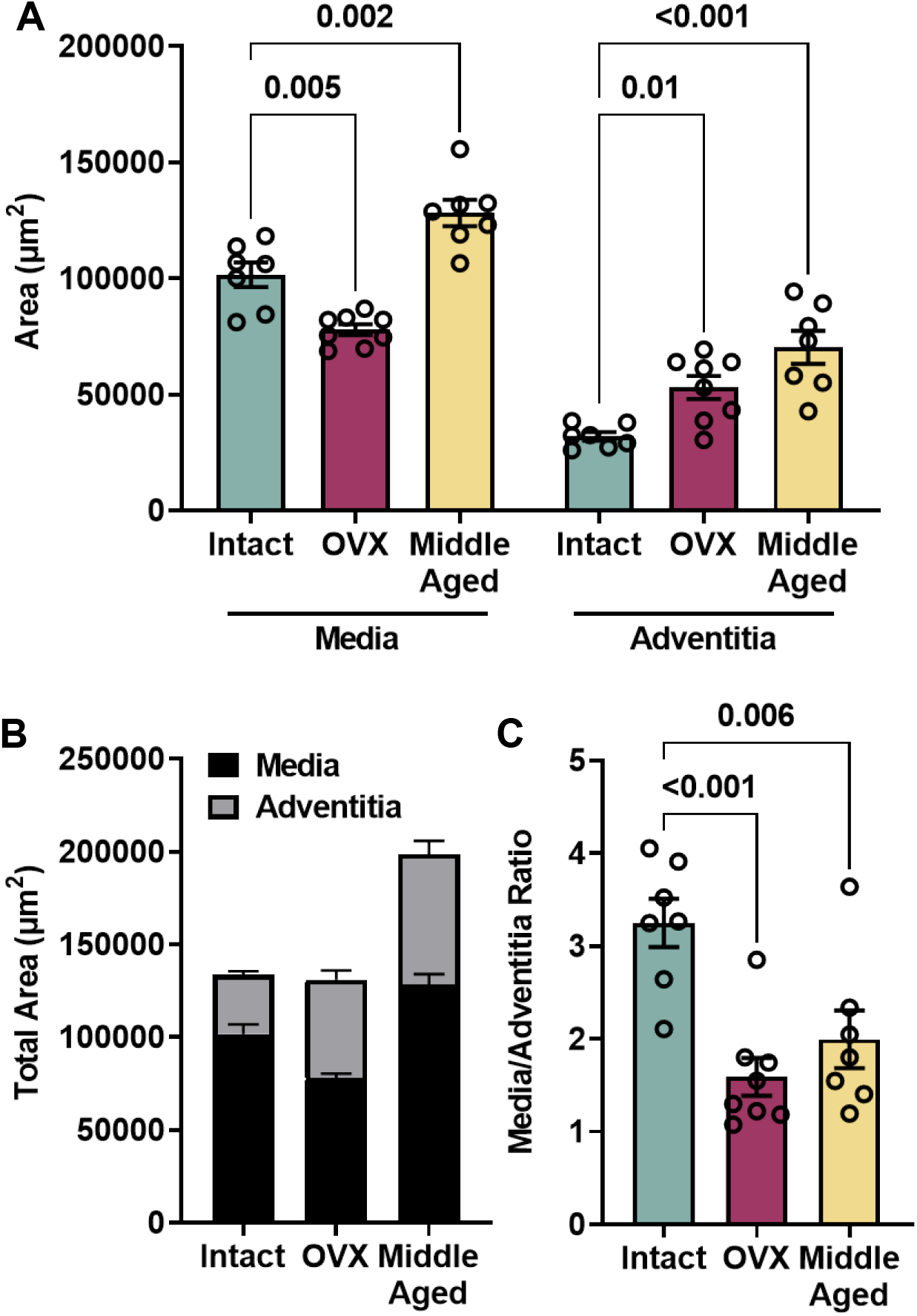
Impact of Aging and OVX on Aortic Wall Composition. Values are mean ± SEM, N=10-11 per group. **(A)** The area of media and adventitia were analyzed separately by one-way ANOVA with Bonferroni’s post-hoc tests, both P<0.001. Medial area was significantly decreased in OVX but increased with aging while adventitial area was increased by both OVX and aging. **(B)** Summed data show the contribution of each layer to overall cross-sectional area. **(C)** The media to adventitia ratio is significantly reduced in both treatment groups, one-way ANOVA, P<0.001.

To identify global changes in gene expression that may contribute to the stiffness in OVX aortas that occurred in the absence of vascular thickening, bulk RNA-sequencing was performed. Gene set enrichment analysis (GSEA) identified 272 gene sets that were significantly enriched in intact controls and 13 gene sets significantly enriched in OVX. The top 10 gene sets for each are shown in Figure 7A. Gene sets enriched in intact control aortas in the Reactome Pathway analysis included three pathways on Notch signaling, smooth muscle contraction, gap junctions, extracellular matrix factors, and interleukin signaling. Gene sets enriched in OVX aortas included multiple pathways related to cellular respiration and mitochondrial function, along with striated muscle contraction. A list of the top 50 genes enriched in each condition are shown as heatmaps (Figure 7B). The impact of OVX on smooth muscle contractile genes was of particular interest, as it is known that arterial stiffening can be due to alterations in vascular smooth muscle cell phenotype. We found that in addition to the Reactome smooth muscle contraction gene set, the much larger KEGG smooth muscle contraction gene set showed significant enrichment in intact aortas (Figure 7C). Based on what has been previously published, we looked at individual values for genes associated with the contractile and synthetic phenotypes (Figure 7D). OVX induced a significant decrease in contractile genes (two-way ANOVA, P<0.002; Figure 7D, left), but surprisingly this was not associated with a concomitant increase in genes associated with the synthetic or proliferative phenotype. In fact, OVX also had an overall effect to suppress proliferative genes (two-way ANOVA, P=0.05; Figure 7D, right). Hence, we validated three genes from the smooth muscle contraction analysis using ddPCR.

**Figure 7.**
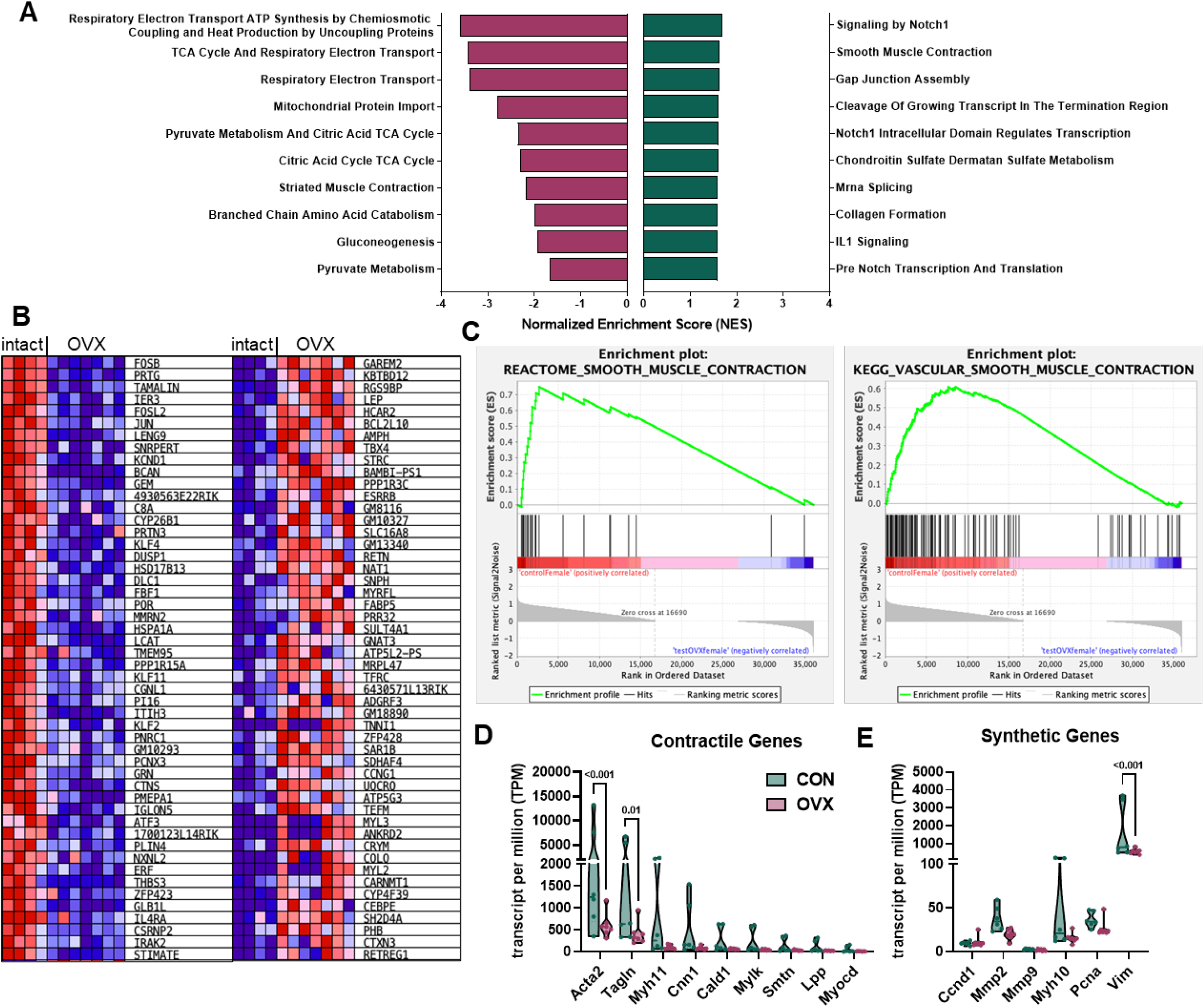
Gene set enrichment (GSEA) analysis of intact and OVX aortas. Bulk RNA-Seq was performed on aortas collected from young adult female mice after OVX for 8 weeks compared to aortas from intact mice (control). **(A)** Reactome enrichment analysis showing the top 10 enriched gene sets in both conditions, all with nominal p value < 0.05. A positive normalized enrichment score (NES) indicates enrichment in intact controls (green), while a negative NES indicates enrichment in OVX (purple). **(B)** Heat map of the top 50 genes for each phenotype. Expression values are represented as a range from red (high expression) to blue (low expression). **(C)** Both Reactome and KEGG gene analysis for gene sets associated with smooth muscle cell contraction show enrichment in intact control aortas. **(D)** Selected contractile genes showed significant downregulation by OVX, two-way ANOVA, P<0.002. **(E)** OVX also decreased expression of synthetic genes, P=0.05.

To determine whether GPER activation reverses OVX-induced arterial stiffening, a separate cohort of mice were OVX at 10 weeks and treated from weeks 18-20 with either vehicle, G-1, or LNS8801. Body weight, heart weight, kidney weight, and uterine weight were not significantly different between any of the groups at the end of the study (Figure 8A-D). Additionally, no changes were detected in systolic blood pressure or pulse pressure due to treatment (Figure 8E-F). OVX significantly increased PWV after 8 weeks, confirming previous data (P<0.0001; Figure 9A). G-1 treatment tended to decrease PWV but did not reach statistical significance (P=0.07), while LNS8801 dramatically reduced PWV to levels observed before OVX (P=0.004; Figure 9B). Neither G-1 nor LNS8801 impacted vascular geometry (Figure 10), which was not surprising considering that there was also no difference in these parameters between control and OVX. The impact of the drug treatments on *ex vivo* material stiffness matched the trends in PWV, with LNS8801 producing a greater rightward shift than G-1 (Figure 11B). Only LNS8801 induced a significant reduction in beta stiffness index (Figure 11C).

**Figure 8.**
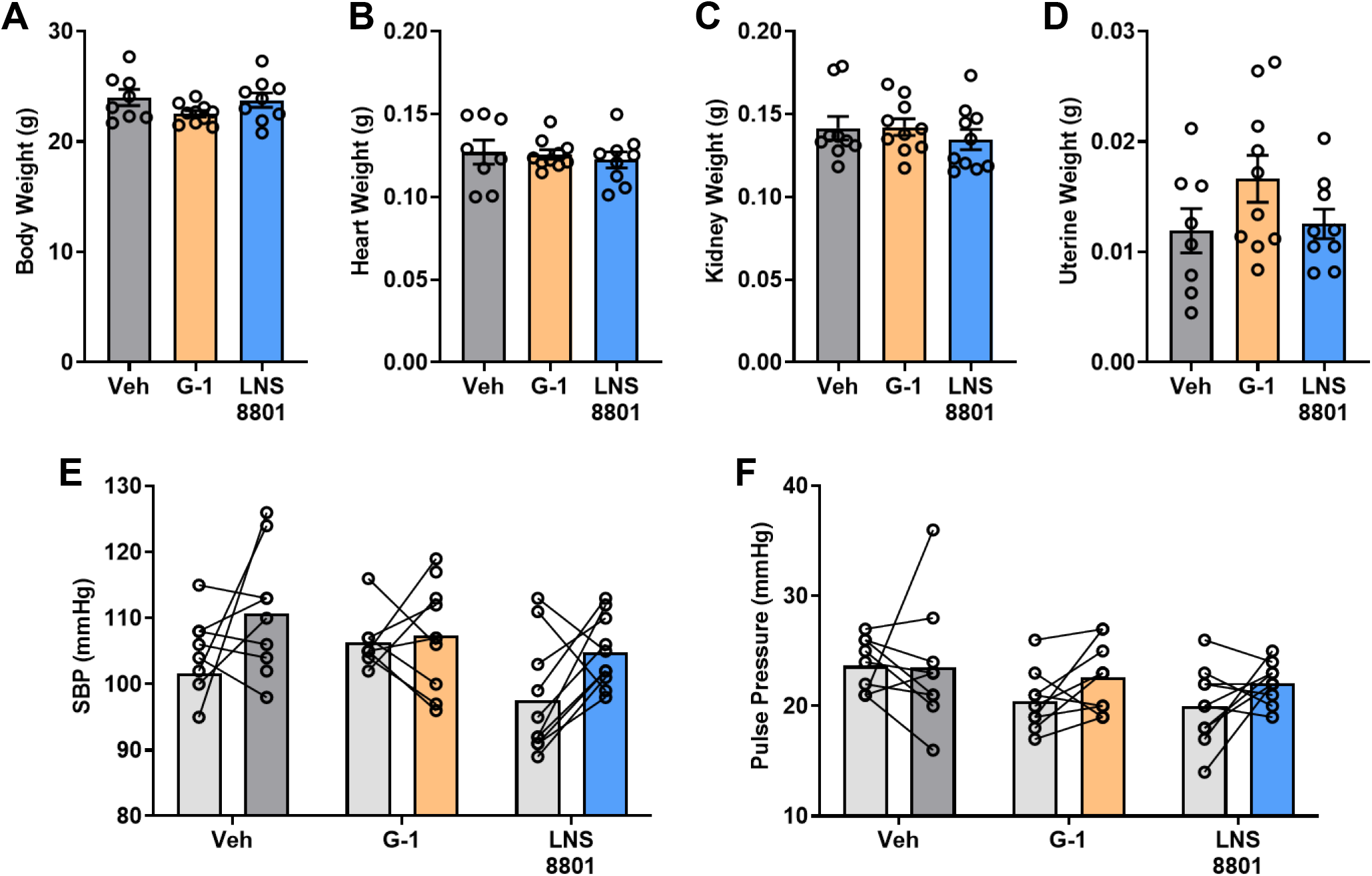
Impact of GPER activation on body weight, tissue weights, and blood pressure. Values are mean ± SEM, N=8-10 per group. No significant differences were found by one-way ANOVA in **(A)** body weight, P=0.15, **(B)** heart weight, P=0.80, **(C)** kidney weight, P=0.63, or **(D)** uterine weight, P=0.17. Repeated measures ANOVA did not find changes in **(E)** systolic blood pressure (SBP) or **(F)** pulse pressure (PP) between pre-treatment (gray bars) to post-treatment or between treatment groups.

**Figure 9.**
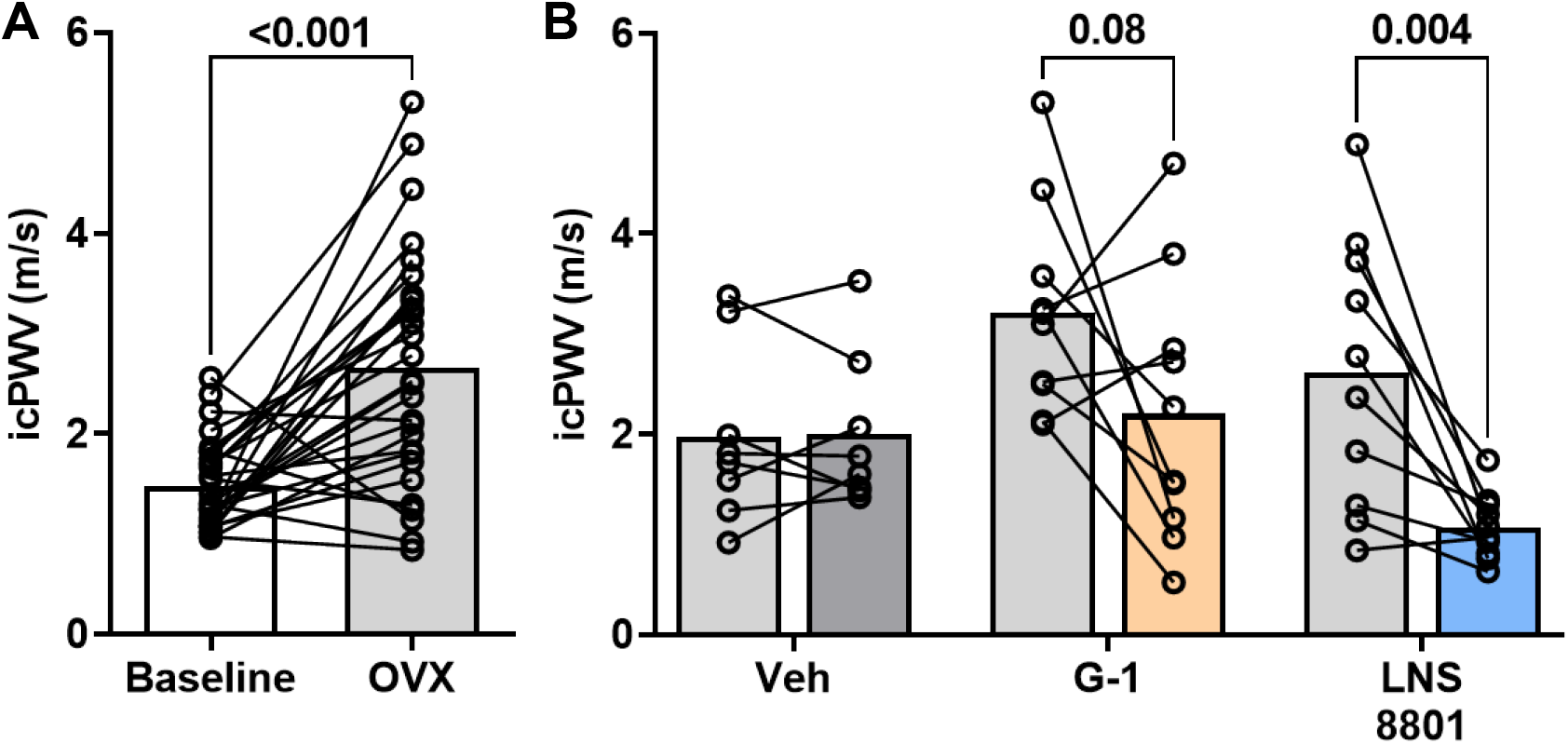
Impact of GPER activation on PWV. Values are mean ± SEM, N=8-10 per group. **(A)** Ovariectomy significantly increased PWV from baseline, paired t-test. **(B)** First bar in each group is PWV 8 weeks post-OVX, and second bar shows values after two-week treatment. Repeated measures ANOVA showed no impact of vehicle (Veh) on intracarotid PWV (icPWV). G-1 treatment caused a downward trend but was not significant, while LNS8801 significantly reduced icPWV.

**Figure 10.**
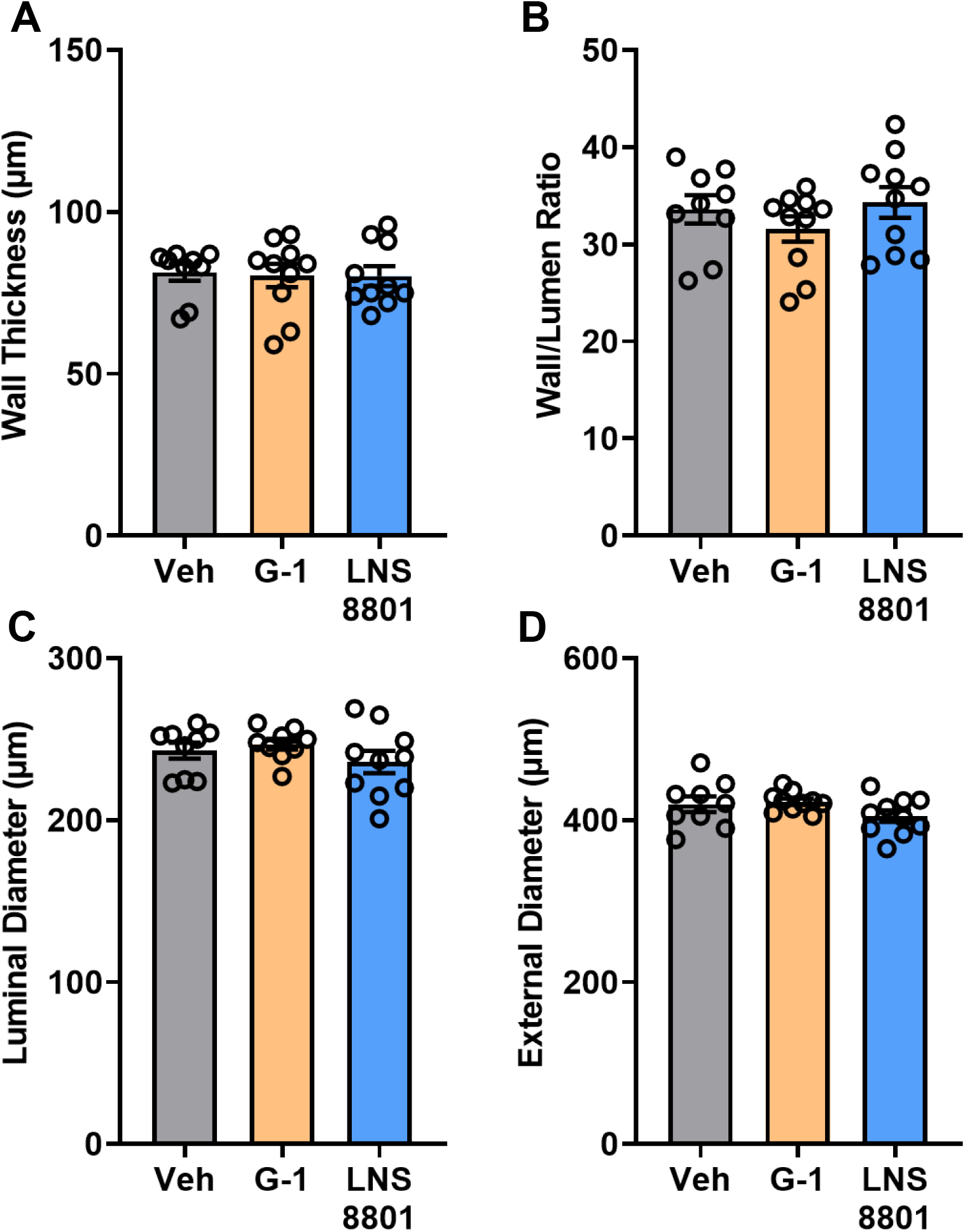
Impact of GPER activation on vascular geometry. Statistical analysis was performed using one-way ANOVA with Bonferroni’s multiple comparisons test shown on the graphs. Values are mean ± SEM, N=9-10 per group. **(A)** Wall thickness, **(B)** wall-to-lumen ratio, **(C)** luminal diameter, and **(D)** external diameter were not different between treatment groups, P>0.05.

**Figure 11.**
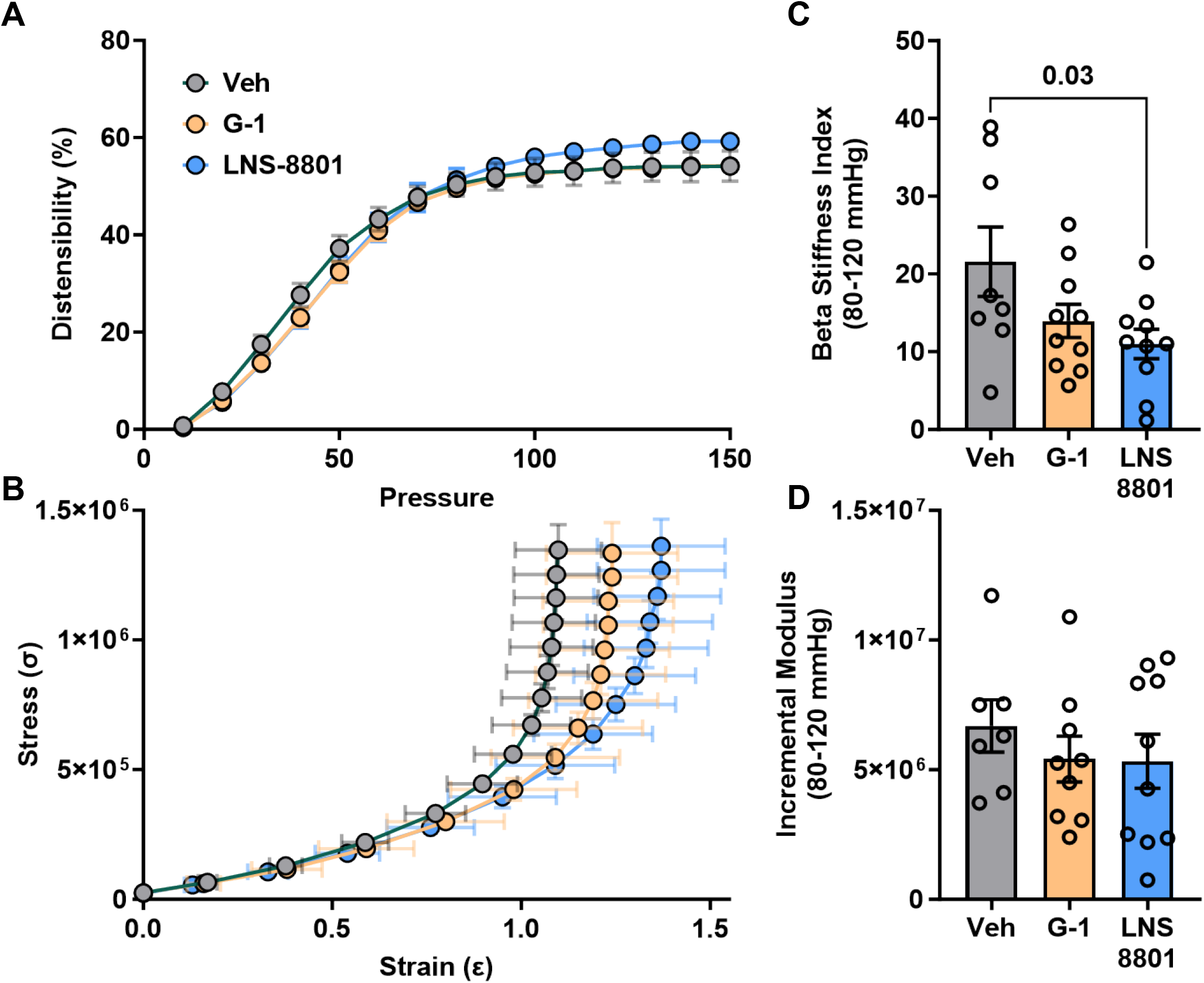
Impact of GPER activation on material stiffness. Values are mean ± SEM, N=8-10 per group. **(A)** Arterial compliance shown as percent distension from baseline showed no significant differences across treatment. Two-way ANOVA, P=0.48. **(B)** A rightward shift in the stress-strain curve was observed for both G-1 and LNS8801 treatment groups. **(C)** Beta stiffness but not **(D)** incremental modulus calculated between 80-120 mmHg showed a decrease in stiffness with LNS-8801. One-way ANOVA, P=0.04 and P=0.60.

Histological analysis showed that G-1 but not LNS8801 increased medial area compared with vehicle and LNS8801 treatment, however, adventitial area was unaffected (Figure 12A). Alcian blue staining for glycosaminoglycans showed no significant impact of treatment (Figure 12B). Since aortic wall composition was not changed, we investigated the impact of LNS8801 on the expression of contractile genes identified in RNAseq analysis. We probed for the three contractile genes with the highest expression in the aorta that were decreased by OVX (Figure 7D). We found that smooth muscle actin (Acta2), transgelin (Tgln), and myosin heavy chain 11 (Myh11) decreased with OVX and were partially restored by two weeks of LNS8801 treatment (Figure 12C). Aortic mRNA expression of GPER was not impacted by OVX or treatment with G-1 or LNS8801 (Figure 12C).

**Figure 12.**
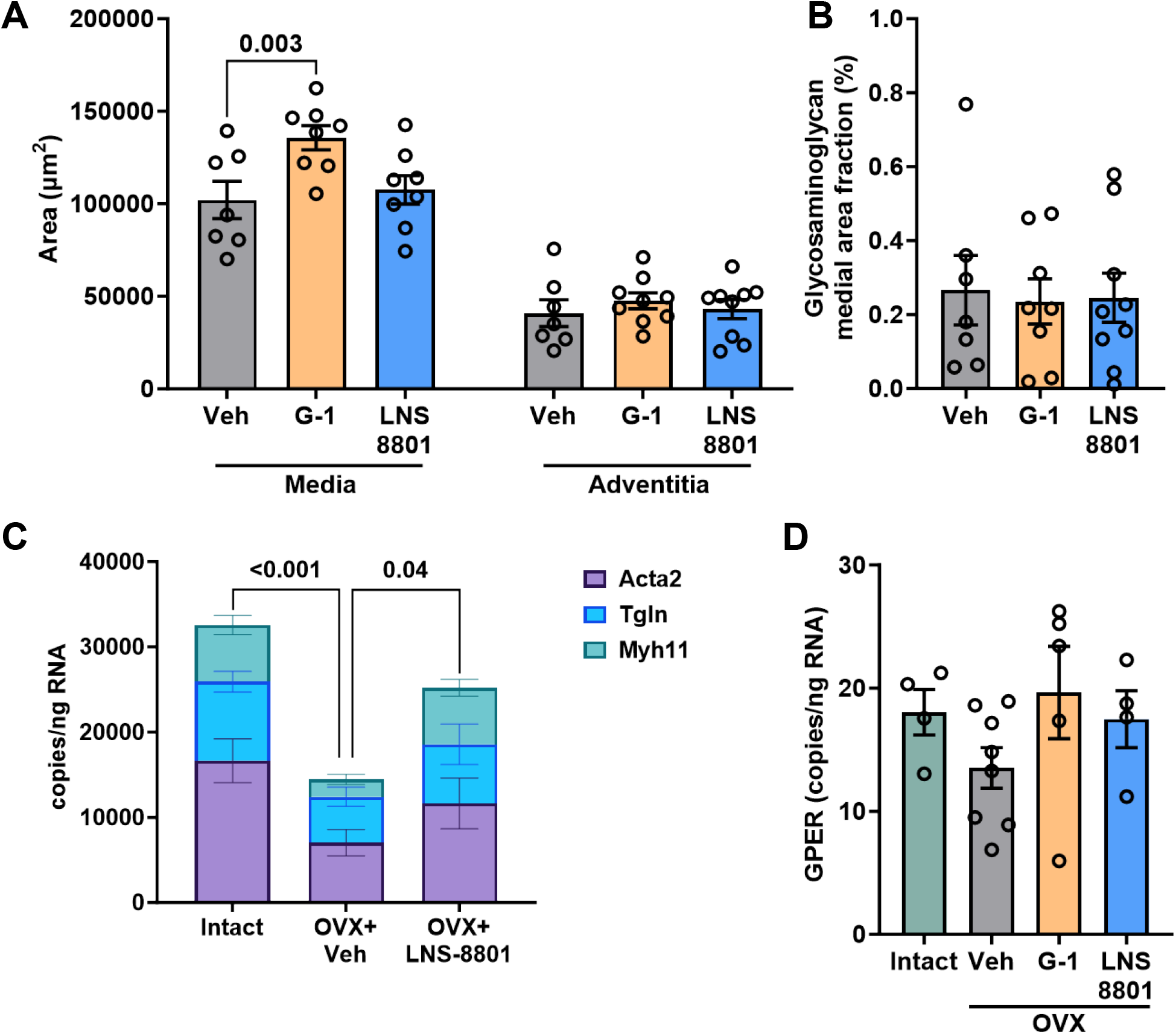
Impact of GPER activation on aortic wall composition and contractile genes. (A) Medial area was changed only in OVX animals treated with G-1. One-way ANOVA, P=0.02. Adventitial area in OVX animals was not different across treatment groups. One-way ANOVA, P=0.82. **(B)** Treatment did not alter glycosaminoglycan content. One-way ANOVA, P=0.96. **(C)** GPER mRNA in aortic tissue was unaffected by OVX or treatment with G-1 or LNS8801. One-way ANOVA, P=0.27. **(D)** Contractile gene expression was different between groups. One-way ANOVA, P<0.001. Multiple comparisons for main effect showed a decrease with OVX (P<0.001) and an increase with LNS8801 (P=0.04).

## DISCUSSION

The current study found that PWV increased to a similar extent in response to either OVX at 10 weeks or aging to 52 weeks, yet the underlying mechanisms were different. Aging was associated with a significant increase in wall thickness due to inward remodeling and an increase in collagen content, while OVX increased material stiffness while decreasing the smooth muscle layer. The loss of the medial layer in OVX mice was associated with a decrease in smooth muscle contractile genes detected by bulk RNA-Seq. Administration of the enantiomerically pure GPER agonist, LNS8801, effectively reversed the impact of OVX on PWV and material stiffness and was associated with partial restoration of contractile genes despite no changes in wall composition. Taken together, these studies show that hormone decline in females is not merely early vascular aging but induces a different mechanism to promote arterial stiffening.

Intracarotid PWV increased in both OVX and middle-aged mice without a significant change in blood pressure. This is consistent with results reported by our lab and other groups showing that PWV is higher in aged mice compared to young mice and occurs prior to measurable changes in BP (15, 20). Our results also mimic recent clinical data showing that menopause is associated with significant increase in arterial stiffening without any change in blood pressure (18). We propose that implementing measurements of PWV may provide opportunities for early intervention against cardiovascular risk in women undergoing the menopausal transition.

The greater wall thickness in aging vessels provided more tissue for the distribution of stress, which explains the lack of change in material stiffness despite an increase in arterial stiffness measured *in vivo* by PWV. The increased wall thickness was due to a decreased luminal diameter and maintained external diameter, thus enhancing the wall to lumen ratio. Our previous study similarly found that material stiffness did not increase in one-year-old female mice, and while there was a trend for increased wall thickness and decreased lumen diameter it did not reach statistical significance (20). Aging to 36 weeks in mice increases wall thickness in coronary arteries, but in this vascular bed an increase in stiffness was also present (36).

In contrast to aging, OVX induced material stiffness that was independent of changes in vascular geometry. The mechanisms promoting arterial stiffening may differ between estrogen loss and aging, but it is also plausible that stiffening precedes changes in structure and a more prolonged estrogen deprivation would eventually promote increased wall thickness. Few animal studies have investigated the impact of ovariectomy alone in the absence of other diseases. One study using C57BL/6J mice found *decreased* strain and Cauchy stress one-month after OVX along with *increased* intimal thickening (37). Another study conducted in coronary arteries shows similar data to the current study, where neither ovariectomy nor estrogen treatment impacts vessel diameter but estrogen increases distensibility (38). These differing results indicate that each vascular bed most likely undergoes multiple remodeling processes after ovariectomy until some level of homeostasis is obtained. Future studies should investigate estrogen loss across more timepoints to fully understand its impact on the vasculature.

Many studies implicate the extracellular matrix in arterial stiffening, yet only the aging group showed an increase in collagen content within the medial layer. However, both the OVX and aging groups displayed an increase in the amount of adventitia surrounding the aorta, which is primarily made of extracellular matrix proteins. We probed for glycosaminoglycans based on our previous finding that aortic remodeling in the rat aorta was associated with changes in this matrix factor and were modulated by GPER activation (28). In contrast, OVX in the current study *decreased* glycosaminoglycan content within the medial layer. *In vitro* studies indicate that both 17β-estradiol (E2) and progesterone reduce collagen and enhance elastin deposition in aortic smooth muscle (2). Similarly, in ovariectomized primates matrix changes in response to an atherosclerotic diet are ameliorated by estrogen treatment (39). Therefore, the impact of estrogen on the extracellular matrix is most likely exacerbated under pathological conditions.

Vascular smooth muscle cells are unique by differentiating and dedifferentiating multiple times throughout their lifetime (40). The dogma on this phenotypic diversity is that mature, healthy VSMC mostly exist in their differentiated or “contractile” state but then dedifferentiate into a “synthetic” phenotype to allow proliferation and damage repair (41). During cardiovascular disease, this plasticity becomes aberrant and smooth muscle cells remain in the synthetic state, allowing proliferation and promoting arterial stiffening, atherosclerosis, and hypertension (42). The switch to this phenotype is characterized by both downregulation of “contractile” proteins such as smooth muscle α-actin, myosin heavy chain, and calponin along with upregulation of “synthetic” markers which include cell cycle and extracellular matrix proteins such as calmodulin and collagen (43). While previous studies indicate that estrogen slows the switch from contractile to synthetic phenotype in cultured VSMC (44) our data are the first to assess the impact of *in vivo* estrogen loss on smooth muscle phenotype and do not support this yin and yang of contractile and synthetic markers. Recent single-cell RNAseq analysis indicates that vascular smooth muscle can exist in at least six different phenotypes (45), and our preliminary data has uncovered another distinct VSMC phenotype in response to estrogen loss.

Estrogen binds genomic estrogen receptors ERα and ERβ as well as the G protein-coupled estrogen receptor (GPER). Previous work shows that vascular remodeling is inhibited by decreasing ERα or increasing GPER expression *in vivo* (46). The potential detrimental impact of ERα was also demonstrated in endothelial cells, where genetic deletion of this receptor is protective against high fat diet-induced arterial stiffening (3). Previous work from our group has demonstrated that selective GPER activation induces many protective cardiovascular effects including decreased BP (24), renal protection (26), and beneficial effects on arterial stiffening and remodeling (28, 33). In the current study, we used an enantiomerically pure GPER agonist developed by Linnaeus Therapeutics (LNS8801), which contains only the enantiomer in G-1 that activates GPER and is currently in clinical trials for treatment of cancer (ClinicalTrials.gov Identifier: NCT04130516). We allowed arterial stiffness to develop after OVX for 8 weeks before initiating treatment with either LNS8801 or the commercially available racemic agonist G-1. We found that LNS8801 produced a robust decrease in PWV in OVX mice and was associated with upregulation of contractile genes within the aortic wall. Additional studies are needed to determine the molecular mechanisms by which GPER impacts contractile gene expression.

Lastly, we showed that intracarotid PWV is a reliable method for detecting arterial stiffness in mice and has the potential to be applied clinically. Since estrogen loss increased stiffness without altering vascular geometry, PWV may be superior to methods such as carotid intima-to-media thickness since it provides information on vascular compliance. Utilization of PWV as a cardiovascular risk factor in perimenopausal women, may allow for earlier intervention in this group and reduce mortality (47). Additional studies are still needed, however, to determine treatment options for patients with significant arterial stiffening.

In conclusion, this study demonstrates that estrogen loss induced a decrease in vascular compliance that presented differently than aging. Additionally, we found that selective GPER activation with LNS8801 reversed the effects of estrogen loss, indicating potential use as a therapy for decreased vascular compliance after menopause. Future studies are needed to determine whether this was a temporal change that will eventually promote increased wall thickness similar to aging. This work has translational relevance for women who undergo early menopause, removal of the ovaries, aromatase inhibition, or other estrogen-suppressing therapies.

## METHODS

### Animals

Adult (20 ± 2 weeks) and middle-aged (52 ± 2 weeks) female C57Bl/6J mice (N= 8-10 per group) were housed in a temperature-controlled vivarium with free access to drinking water and standard chow and a standard alternating 12-hour dark to light schedule. Mice were euthanized by exposure to isoflurane and secondary exsanguination.

### Ovariectomy (OVX)

OVX was performed under isoflurane anesthesia following sterile technique. Mice were administered 2 mg/kg of meloxicam subcutaneously for pain management and monitored for successful recovery. Uterine horns were ligated with non-absorbable silk suture before removal of the ovaries. The internal muscle layer was sutured, and the outer layer of skin was closed with staples. Successful OVX was confirmed by uterine weight after euthanasia.

### Blood Pressure

Tail-cuff plethysmography was used to measure blood pressure in awake mice around using the Kent Scientific CODA® Noninvasive Blood Pressure System. Measurements were taken for 5 days, including 2 days of training, at approximately the same time each day to avoid differences due to circadian rhythms.

### Pulse Wave Velocity

Intracarotid PWV (icPWV) and aortic PWV (aPWV) were measured using the VisualSonics Vevo® 1100 High-Frequency Ultrasound System as previously described (19, 20). Briefly, isoflurane anesthesia was induced and maintained via nose cone in the supine position on a heated EKG platform. The right carotid artery was imaged in Doppler mode both distal to the aortic arch and proximal to the carotid bifurcation, followed by imaging of the abdominal aorta. PWV analysis was conducted by a different, blinded investigator using VisualSonics Vevo LAB software (v.5.7.0) with PWV measurement package.

### Passive Biaxial Pressure Myography

Carotid arteries were excised and cannulated on 27-gauge needles using 9-0 black mononylon for biaxial pressure myography in Hank’s balanced salt solution (HBSS) as previously described (19, 20). Briefly, carotids underwent standardized testing to determine the estimated in vivo length which was confirmed where the axial force was nearly constant over a pressure of 10-150 mmHg. Next, vessels underwent three cycles of pressure diameter testing where pressure was applied to the lumen of the vessel and increased from 10-150 mmHg then decreased back to 10 mmHg while outer diameter was recorded.

### Histology

Aortas from female mice were formalin-fixed, paraffin-embedded, and placed on microscope slides. Verhoeff-Van Gieson (VVG) stain was used to quantify elastin, smooth muscle, and collagen content, while Alcian blue stained for glycosaminoglycans. Analysis was performed using Adobe Photoshop (v.24.5.0). Data analysis was performed by a different investigator who was blinded.

### Bulk RNA-Sequencing

Bulk RNA-Sequencing was performed using whole aortas of adult female intact (n=4) and OVX (n=7) mice. Aortas were stored in RNAlater (Thermo Fisher) at 4°C for 24 hours, then moved to −20°C for long term storage. Total RNA samples were isolated using RNeasy Plus Mini Kit (Qiagen). Nanodrop was used for RNA quantity followed by QA/QC using Agilent 2100 Bioanalyzer. Nextseq mid output (130M clusters) and single read 150 cycles kit was used. Sequences in paired-end fastq files, 35-76bp long, were assessed for quality using FASTQC (v.0.11.7). Reads were pseudoaligned to mouse reference transcriptome (mm10) and expression quantified using Kallisto (v0.46.0). Principal component analysis was used to exclude samples that were not clustering with their respective group. Differential expression was performed using DESeq2 (v.1.38.3). Expression profiles were identified using Gene Set Enrichment Analysis (GSEA, v.4.0.3). Differential splicing analysis was performed using rMATS-turbo (v.4.1.1).

### Drug Treatment

A second cohort of mice underwent OVX at 10 weeks and were randomized at 18 weeks to receive two-week treatment with either vehicle, the racemic GPER agonist G-1 (Cayman Chemical), or the purified active enantiomer LNS8801 (Linnaeus Therapeutics). The dose of G-1 and LNS8801 was 400 µg/kg/day based on our previous publication (24). G-1 and LNS8801 were first dissolved in dimethyl sulfoxide then combined with an equal volume of 30% ethanol at 37°C to prevent precipitation. Drug or vehicle was added to osmotic minipumps (Alzet Model 1002), primed in sterile saline for 24 h at 37°C, and implanted subcutaneously under isoflurane anesthesia.

### Droplet digital PCR (ddPCR)

RNA was extracted from aortas using the RNeasy Plus Mini Kit (Qiagen 74136), and ddPCR was conducted as previously described (48, 49) using the following validated mouse primers were obtained from Bio-Rad: Acta2 (dMmuCPE5117282), Tagln (dMmuCPE5113593), and Myh11 (dMmuCPE5116710).

### Statistical Analysis

All data are expressed as mean ± SEM. Data were analyzed using GraphPad Prism (v.9.5.1). Individual replicates are represented as symbols on graphs and bars represent the mean values for each group. Statistical tests and results are presented in each figure legend, and differences were considered significant at P<0.05.

### Study Approval

Animal treatments and procedures were in accordance with the National Institutes of Health Guide for the Care and Use of Laboratory Animals and were approved by the Tulane University Institutional Animal Care and Use Committee. All mice were maintained and aged in the Tulane University animal facility.

## Acknowledgements

Funding was provided by the National Institutes of Health (HL155841 to B.O.O., HL133619 and AG071746 to S.H.L) and the American Heart Association (559076 to B.V. and 559082 to I.K.). Assistance with RNA-Seq analysis was provided by the Tulane Cancer Center Cancer Crusaders Next Generation Sequence Analysis Core supported by the National Institutes of Health National Cancer Institute, Award Number P01CA214091. The content is solely the responsibility of the authors and does not necessarily represent the official views of the National Institutes of Health.

## Data Availability

Data sets generated for this study are available in Harvard Dataverse (*link added upon publication*).

## Authors Contributions

I.K. and S.H.L. designed the research studies, analyzed the data, and wrote the manuscript. I.K., A.M., T.W., B.V., S.B., A.I.S., C.R., Z.D., A.H., and B.O.O. conductedthe experiments and acquired data. C.A.N. provided reagent. All authors read and edited the manuscript.

